# Self beyond the body: task-relevant distal cues modulate performance and body ownership

**DOI:** 10.1101/361022

**Authors:** Klaudia Grechuta, Laura Ulysse, Belén Rubio Ballester, Paul F.M.J. Verschure

**Affiliations:** Department of Information and Communication Technologies, Universitat Pompeu Fabra (UPF), 08-018 Barcelona, Spain; Institute for Bioengineering of Catalonia (IBEC), The Barcelona Institute of Science and Technology (BIST), 08-028 Barcelona, Spain; Pompeu Fabra University, Center for Brain and Cognition, Computational Neuroscience Group, Department of Information and Communication Technologies, 08-018 Barcelona Spain; Catalan Institution for Research and Advanced Studies (ICREA), 08-010 Barcelona Spain

**Author notes:** These authors equally contributed to this study (Klaudia Grechuta, Laura Ulysse). moved to (Laura Ulysse).

## Abstract

The understanding of Body Ownership (BO) largely relies on the Rubber Hand Illusion (RHI) where synchronous stroking of real and Rubber Hands (RH) leads to an illusion of ownership of RH provided physical, anatomical, postural and spatial plausibility of the two body-parts. RHI also occurs during visuomotor synchrony, in particular, when the visual feedback of virtual arm movements follows the trajectory of the instantiated motor command. Hence BO seems to result from a bottom-up integration of afferent and efferent proximal multisensory evidence, and top-down prediction of both externally and self-generated signals, which occurs when the predictions about upcoming sensory signals are accurate. In motor control, the differential processing of predicted and actual sensory consequences of self-generated actions is addressed by, the so-called, Forward Model (FM). Based on an efference copy or corollary discharge, FM issues predictions about the sensory consequences of motor commands and compares them with the actual outcome. The discrepancies (Sensory Prediction Errors, SPEs) are used to correct the action on the consecutive trial and provide new estimates of the current state of the body and the environment. Here, we propose that BO might be computed by FMs, and therefore, it might depend on their consistency, specifically, in contexts where the sensory feedback is self-generated. Crucially, to reduce SPE, FMs integrate both proximal (proprioceptive) and distal (vision, audition) sensory cues relevant to the task. Thus, if BO depends on the consistency of FMs, it would be compromised by the incongruency of not only proximal but also distal cues. To test our hypothesis, we devised an embodied VR-based task where action outcomes were signaled by distinct auditory cues. By manipulating the cues with respect to their spatiotemporal congruency and valence, we show that distal feedback which violates predictions about action outcomes compromises both BO and performance. These results demonstrate that BO is influenced by not only efferent and afferent cues which pertain to the body itself but also those arising outside of the body and suggest that in goal-oriented tasks BO might result from a computation of FM.

## Introduction

Humans and other species simultaneously acquire and integrate both self-generated (reafferent) and externally-generated (exafferent) information through different sensory channels [1]. Imagine performing a goal-oriented task such as an Air Hockey where the objective is to score points by hitting a puck into the goal. During the task, the brain continuously stores and updates both (1) action-driven proximal (ie. proprioceptive) and distal (ie. visual or auditory) sensory cues relevant to the task such as the kinematics of the arm or the location of the puck [2, 3], and (2) action-independent signals driven by external contingencies such as wind [4]. Thus the ability of the nervous system to determine the source of a given sensation and the boundaries of an embodied self, commonly referred to as the sense of Body Ownership (BO) [5, 6], is fundamental in adaptive goal-oriented behavior [7, 8]. However, mechanisms driving BO in action contexts, which require manipulation of the environment and therefore integration of distal cues (ie. Air Hockey) remain elusive.

Our understanding of BO largely relies on, the so-called, Rubber Hand Illusion (RHI) paradigm where the participants experience the illusion of ownership towards a Rubber Hand (RH) during externally-generated synchronous, but not asynchronous, stroking of both real and artificial hands [5]. The illusion generalizes to distinct body-parts including fingers, face or a full body [9–11]. Interestingly, further studies show that while BO strongly depends on the spatiotemporal alignment of the exteroceptive inputs, it also requires physical, anatomical, postural and spatial plausibility of the two body-parts [3, 12–14]. Hence, this empirical framework proposes the notion that BO is a sensory state which does not require motor commands and relies on two complementary processes. In particular, (1) a bottom-up accumulation and integration of afferent, proximal, multisensory (proprioceptive, tactile) evidence, and (2) top-down comparison between the novel sensory stimuli (RH) and self-specific priors based on the internal model of body representation [12, 15, 16].

Interestingly, a body of recent studies extends the traditional view on the mechanisms driving BO by demonstrating that it does not exclusively stem from the integration of afferent sensory signals. Specifically, it has been shown that BO can also emerge in a top-down manner as (1) a consequence of pure expectation of correlated exafference and (2) an accurate prediction of sensory consequences following self-generated movement. The former finding comes from the study of Ferri and colleagues [17], were subjects experience BO when observing the experimenter’s hand approaching the RH and anticipating the sensory event in the absence of actual touch. The latter, in turn, is supported by the results from Virtual Reality (VR)- based experiments investigating the effects of visuomotor (a)synchrony on BO during goal-oriented behavior [18–20]. Here, the authors report high BO in the condition where movements of the real and virtual hands are spatiotemporally congruent. In particular, when the actual sensory feedback (visual and proprioceptive) of the motor commands matches the expected one. This evidence further highlights the crucial role of the prediction-driven top-down processes in the modulation of BO [16, 21, 22].

In the contemporary theoretical framework of motor control, the differential processing of the predicted and the actual sensory consequences of self-generated actions is addressed by, the so-called, internal Forward Model (FM) [23–25] which issues predictions about sensory consequences of current motor commands. According to this interpretation, in the Air Hockey task at every trial, the motor system prepares and generates those commands which are most likely to cause the desired outcome. Simultaneously, an Efference Copy (EC) [26] or a Corollary Discharge (CD) [27, 28] of these motor signals is sent to the FM. Based on a history of sensorimotor contingencies and the current EC [29], the FM simulates and predicts both the proximal (proprioceptive feedback from the upper limb muscles) and distal (visual or auditory feedback of the puck hitting the goal spatiotemporally aligned with the trajectory) sensory consequences of the efferent movement [7, 30]. Upon action completion, the predicted and the actual information from the sensory systems are compared and the discrepancies constitute error signals, the so-called Sensory Prediction Errors (SPE) [31]. FM integrates them to (1) correct the motor command on the consecutive trial to maximize future performance [31, 32], and (2) provide new estimates of the current state of the body and the environment allowing self-recognition [16, 21, 22, 31, 33].

Grounded in the discussed neurophysiological evidence about the nature of the internal FMs and their functions in goal-oriented behavior as well as the novel insights about BO, here, we propose that BO might be a product of the computation of FMs and therefore it might depend on their consistency, specifically, in action contexts where sensory signals are self-generated. Crucially, FMs are not limited to the bodily feedback exclusively, but rather they integrate across both proximal and distal sensory predictions which pertain to the interactions of an agent within an environment in a goal-oriented manner [30]. We will refer to these signals as Task-Relevant Proximal Cues (TRPC) and Task-Relevant Distal Cues (TRDC), respectively. For instance, the visual and auditory feedback of the puck hitting the goal (TRDC) is spatiotemporally aligned with its trajectory that depends on the direction of the arm movement. In case the actual location of the sound of the puck hitting the goal does not correspond to the efference copy or corollary discharge, it would reflect on errors of FMs [23, 24, 29, 34, 35] and, according to our hypothesis, errors in BO. To test our prediction, we devise an embodied VR-based goal-oriented task where action outcomes are signaled by distinct auditory TRDC. We manipulate the cues with respect to their spatiotemporal congruency and valence and compare BO and performance across two experimental conditions, where the signals are either congruent or incongruent. Our results demonstrate, for the first time, that TRDCs which violate predictions about the consequences of action-driven outcomes compromise both BO and performance.

## Materials and Methods

### Participants

After providing written informed consent, sixteen healthy participants were recruited for the study, eight males (mean age 24.0 *±* 2.65) and eight females (mean age 22.64 *±* 2.25). All subjects were right-handed (handedness assessed using Edinburgh Handedness Inventory) [36], had normal or corrected-to-normal vision and reported normal hearing. They were pseudorandomly assigned to two experimental groups following a between-subjects design, which prevented habituation to the ownership measures, distal auditory cues, manipulations and fatigue. We used stratified randomization to balance the conditions in terms of age, gender and previous experience with VR. All participants were blind to the purpose of the study. The experimental procedures were previously approved by the ethical committee of the University of Pompeu Fabra (Barcelona, Spain).

### Task: VR-based Air Hockey

The experimental setup (Fig 1 A) comprised a Personal Computer (PC), a motion detection system (Kinect, Microsoft, Seattle), a Head Mounted Display (HMD, HTC Vive, www.vive.com) and headphones. Similar to others [19, 37], here we used VR method as a tool to study the modulation of BO. The protocol was integrated within the Virtual Environment (VE) of the Rehabilitation Gaming System (RGS) [38]. During the experiment, while seated at a table, participants were required to complete a goal-oriented task that consisted in hitting a virtual puck into the goal (air hockey, Fig 1 A, B1). Throughout the experiment, participants’ arm movements were continuously tracked and mapped onto the avatar’s arm, such that the subjects interacted with the VE by making planar, horizontal movements over a tabletop (Fig 1 A, B). To prevent repetitive movements, at the beginning of every trial, the puck pseudorandomly appeared in one of the three Starting Positions (SP; left, center, right) (Fig 1 B2). The frequency of appearance of every SP was uniformly distributed within every experimental session. Participants received instructions to place their hand in an indicated SP and to execute the movement to hit the puck when the color of the SP changed to green (”go” signal). Each trial consisted of one “hit” which could end in either a success (the puck enters the goal) or a failure (the puck hits one of the three walls). At the end of every trial, participants were to place their left hand back on the SP. The Experimental Block (EB), in both conditions, consisted of 150 trials preceded by 20 trials of Training (TB) (Fig 1 D) and followed by a Threatening Event (TE). TE served to measure autonomous responses to an unexpected threat (BO measure, Fig 1 C) [39]. Overall, the task had an approximate duration of 20 minutes.

### Task-Relevant Distal Cues Manipulations

The task included Task-Relevant Distal Cues (TRDC) in a form of auditory feedback which was triggered as a consequence of every interaction of the puck with the environment. In particular, at the end of every trial, an auditory cue indicated a success (positive sound) or a failure (negative sound). To study whether TRDC influence BO and performance, we manipulated the congruency of the auditory stimuli in three domains (Fig 1 E) — temporal: the time of the cue was synchronized with the time of the hit; spatial: the cue originated from the location of the hit; and semantic: the valence of the cue reflected performance. The TRDC were manipulated in two experimental conditions including congruent (”C”) and incongruent (”I”). In the training block and “C”, TRDC were always congruent such that they occurred at the time of the hit, at the location of the hit, and they reflected performance (success or failure). In “I”, TRDC were always incongruent, namely, they were anticipated or delayed (temporal domain), they originated in the opposite location of the hit (spatial domain) or they did not reflect performance (semantic domain). All manipulations were pseudorandomly distributed and counterbalanced within each session to counteract order effects. Importantly, Task-Relevant Proximal Cues (TRPC) such as the visual feedback on arm movements remained congruent in both conditions.

### Measures

#### Motor control

We used three measures to quantify performance: scores, directional error, and Reaction Times (RTs). Scores were calculated as the percentage of successful trials (the puck enters the gate), while the directional error equaled the absolute angular deviation from the straight line between the starting position of the puck (left, central or right) and the center of the gate (Fig 1 B3). We computed RTs as time intervals between the appearance of the puck and action initiation. Since the task did not impose a time limit, we expected neither significant differences in RTs between conditions nor speed-accuracy trade-offs. We predicted, however, that the auditory TRDC manipulations in “I” might alter scores and directional accuracy as compared to “C”.

**Figure 1.**
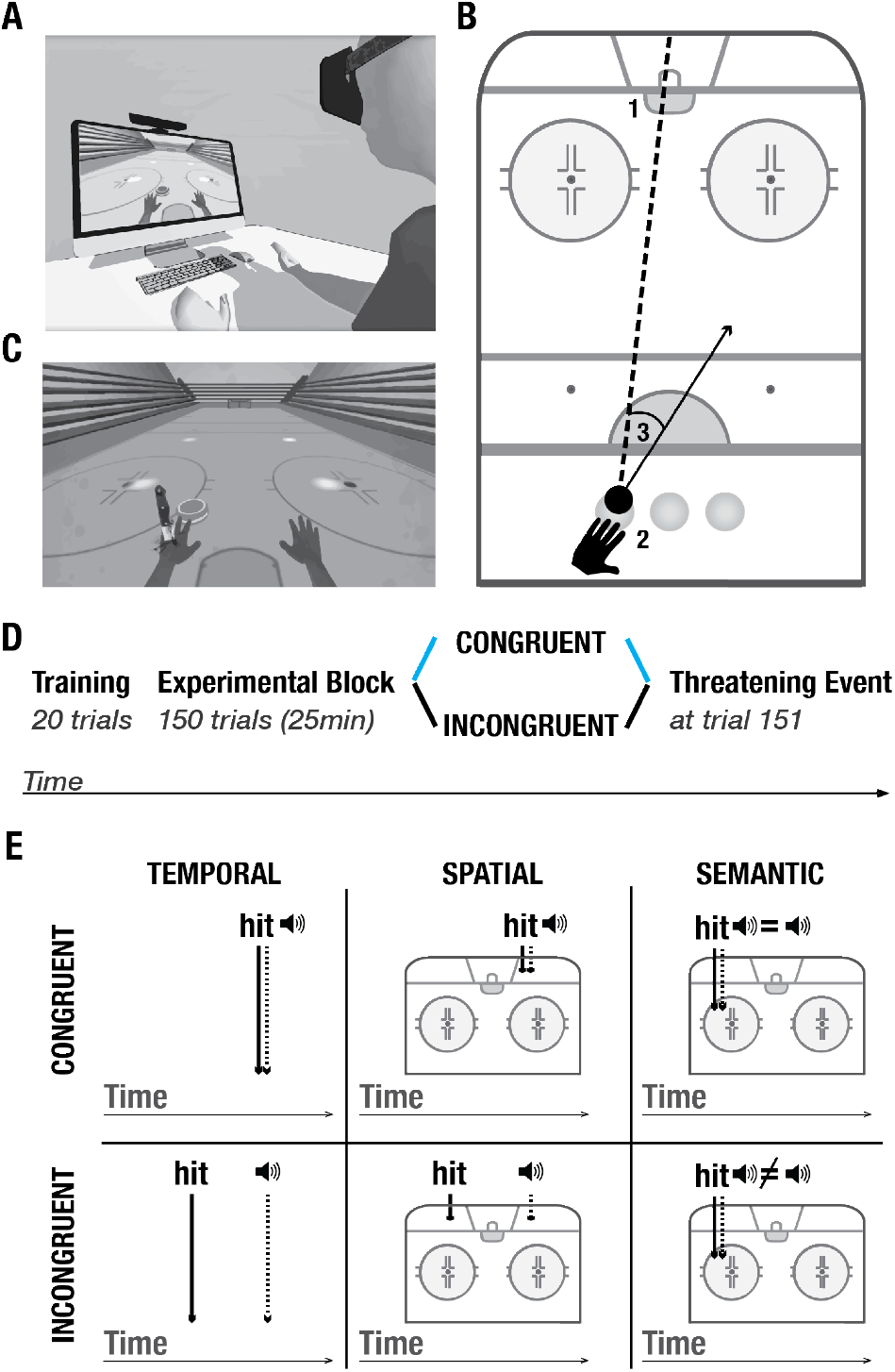
Experimental setup and protocol. (A) Experimental setup. (B). Task. 1- goal. 2- three starting positions. 3- example of a directional error, calculated as the difference between the actual direction vector and a straight line between the position of the puck and the goal. (C) Threatening event. (D) Experimental protocol. All participants underwent the training block. In the experimental block, they were randomly split into two conditions: Congruent “C” (blue), and Incongruent “I” (black). At trial 151, all participants went through the threatening event which served to measure GSR responses. The same color-code (”C”- blue, “I”- black) is used throughout the manuscript. (E) TRDC auditory manipulations- temporal, spatial and semantic. Upper panel: congruent condition; lower panel: incongruent condition.

#### Body ownership

##### Galvanic Skin Response (GSR)

At the end of every experimental session, in both conditions, we introduced a TE (a knife falling down to stab the palm of the virtual hand, Fig 1 C) to quantify autonomous, physiological responses to an unexpected threat [39]. GSR was recorded and stored throughout the experiment. For the analysis, we calculated the mean and the standard deviation of the integral of baseline-subtracted GSR signal per condition in a non-overlapping time windows of 9s. In particular, we expected an increased GSR following the threatening stimulus in “C” as compared to “I”.

##### Proprioceptive drift (PD)

Prior to and upon completion of the experiment, all the subjects completed the PD task which followed a standard technique, see for instance [19]. Specifically, they were asked to point to the location of the tip of their left index finger with the right index finger with no visual feedback available. The error in pointing [12] was computed as the distance between the two locations (the actual location of the tip of the left index finger and the pointing location) and measured in centimeters. We subtracted baseline responses from post-experimental errors for each participant. We expected stronger proprioceptive recalibration, and therefore, higher pointing errors in “C” as compared to “I”.

##### Self-report (S-R)

At the end of every session, all participants completed a questionnaire which evaluated the subjective perception of BO and agency, adapted from previous studies [5, 20, 40]. The entire self-report consisted of twelve items, six per domain (ownership, agency), three of which were related to the experience of ownership and agency respectively, while the remaining served as controls. To counteract order effects, the sequence of the questions was randomized across all the subjects.

## Results

To test our hypothesis that action-driven distal cues which pertain to the task contribute to Body Ownership (BO), we used a VR-based experimental setup (Fig 1 A, B) where subjects were to complete a goal-oriented task by performing horizontal (planar) movements, and manipulated the congruency of TRDC action outcomes (Fig 1 E). The experimental protocol (Fig 1 D) consisted of three phases: Training Block (TB), (2) Experimental Block (EB) in either congruent (”C”) or incongruent (”I”) condition, and (3) Threatening Event (TE) (Fig 1 D, C). To quantify BO, for each experimental session, we measured Proprioceptive Drift (PD), recorded Galvanic Skin Response (GSR) to an unexpected threat, and collected self-reports. To compute performance, we measured scores, directional error, and Reaction Times (RTs). For the analysis, we used t-tests and calculated Cohen’s d to evaluate differences between conditions and the associated effect sizes.

### Motor Control

Firstly, our results showed that the normalized performance-scores (proportion of successful trials) were significantly higher in “C” (*µ* = 0.35, *SD* = 0.47) than in “I” (*µ* = 0.17, *SD* = 0.38), (*t* = 8.89, *p <* 0.001, *d* = 0.42) (Fig 2 A). To explore the effects of the congruency of TRDC on performance, we compared both conditions in terms of directional errors (Fig 2 B). In particular, a T-test indicated that the errors were significantly higher in “I” (*µ* = 6.42, *SD* = 4.52) than in “C” (*µ* = 3.30, *SD* = 2.01), (*t* = *−*19.52, *p <* 0.001, *d* = 0.89) (Fig 2 C). To further investigate the relationship between the quality of the TRDC and performance, we averaged and compared the directional errors following the three types of auditory manipulations (Fig 2 D). Interestingly, we found no difference between the distinct TRDC including spatial (*µ* = 10.17, *SD* = 13.33), temporal (*µ* = 7.99, *SD* = 9.75) and semantic (*µ* = 7.22, *SD* = 7.23) cues. Specifically, a Kruskal-Wallis test indicated that all manipulations had the same significant effect on BO (*KW, p* = 0.39). In addition, we observed that the congruency of the TRDC had no significant effect on the averaged RTs when comparing “I” (*µ* = 0.48, *SD* = 0.05) with “C” (*µ* = 0.51, *SD* = 0.01), *p* = 0.46 (Fig 2 E).

**Figure 2.**
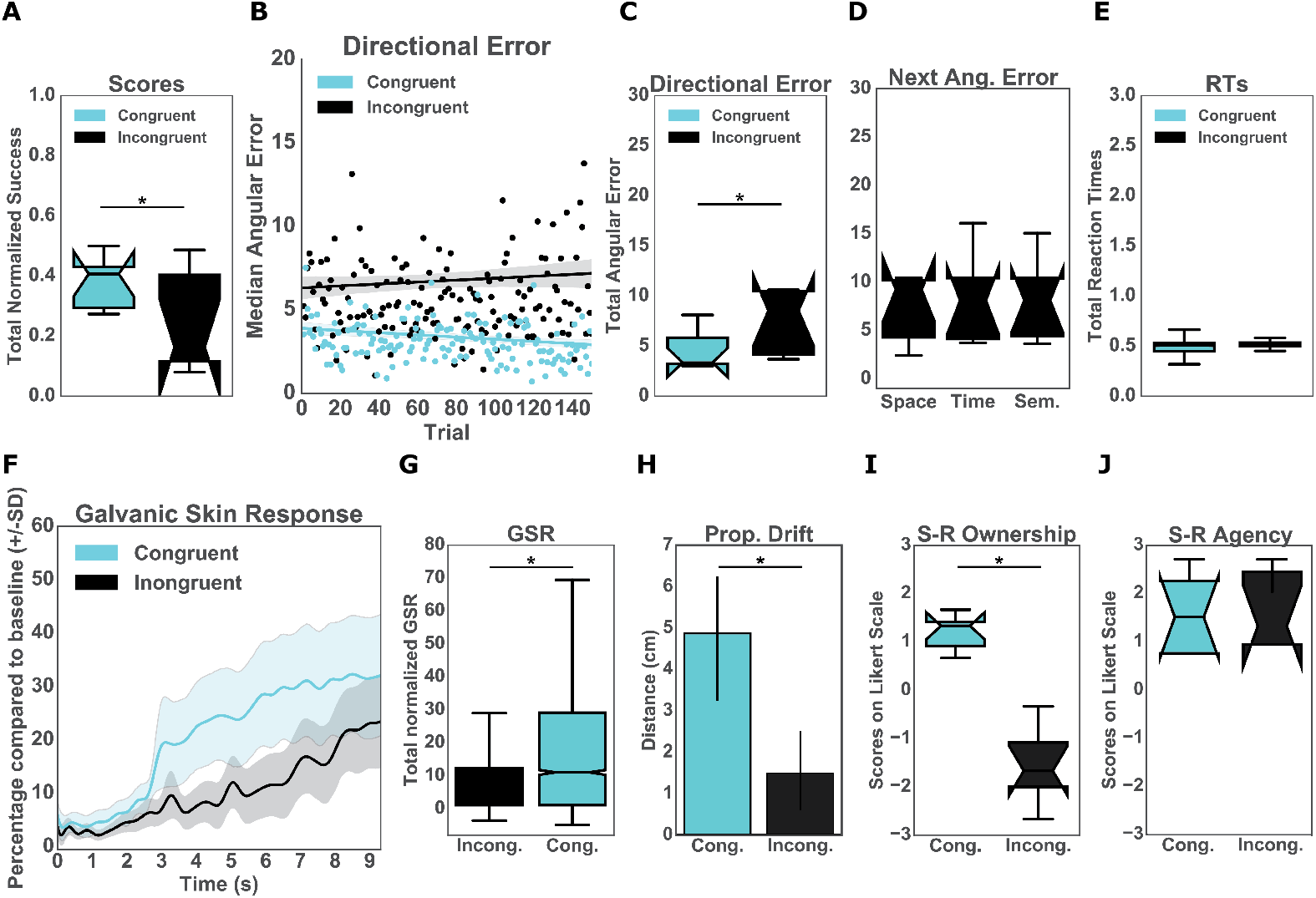
*Upper panel: Performance*. (A) Normalized percentage of successful trials per group. (B) Median directional error per trial over the experimental block (N=150) split per conditions (C) Total directional error from all the trials per subject per condition. (D) The effects of the three auditory manipulations (spatial, temporal and semantic) on the mean directional error on the consecutive trials. (E) Mean reaction times from all trials per condition. *Lower panel: Body Ownership*. (F) Galvanic Skin Response (GSR). The sampling rate for the GSR signal was 60 Hz. Accordingly, the data was run through a low-pass filter with a cut-off frequency of 3 Hz. The plot represents the mean GSR and the associated standard deviation for all participants in a time window of 9s, split per condition. The threatening event happened at time 0. (G) Mean GSR from 9s seconds post threatening event. (H) Proprioceptive Drift (PD). Results of the difference between pre- and post test calculated in centimeters per condition. (I) Score from self-reported (SR) experience of body-ownership per group. Scores above 0 indicate BO. (J) Score from self-reported (SR) experience of agency per group. Scores above 0 indicate the experience of agency.

**Figure 3.**
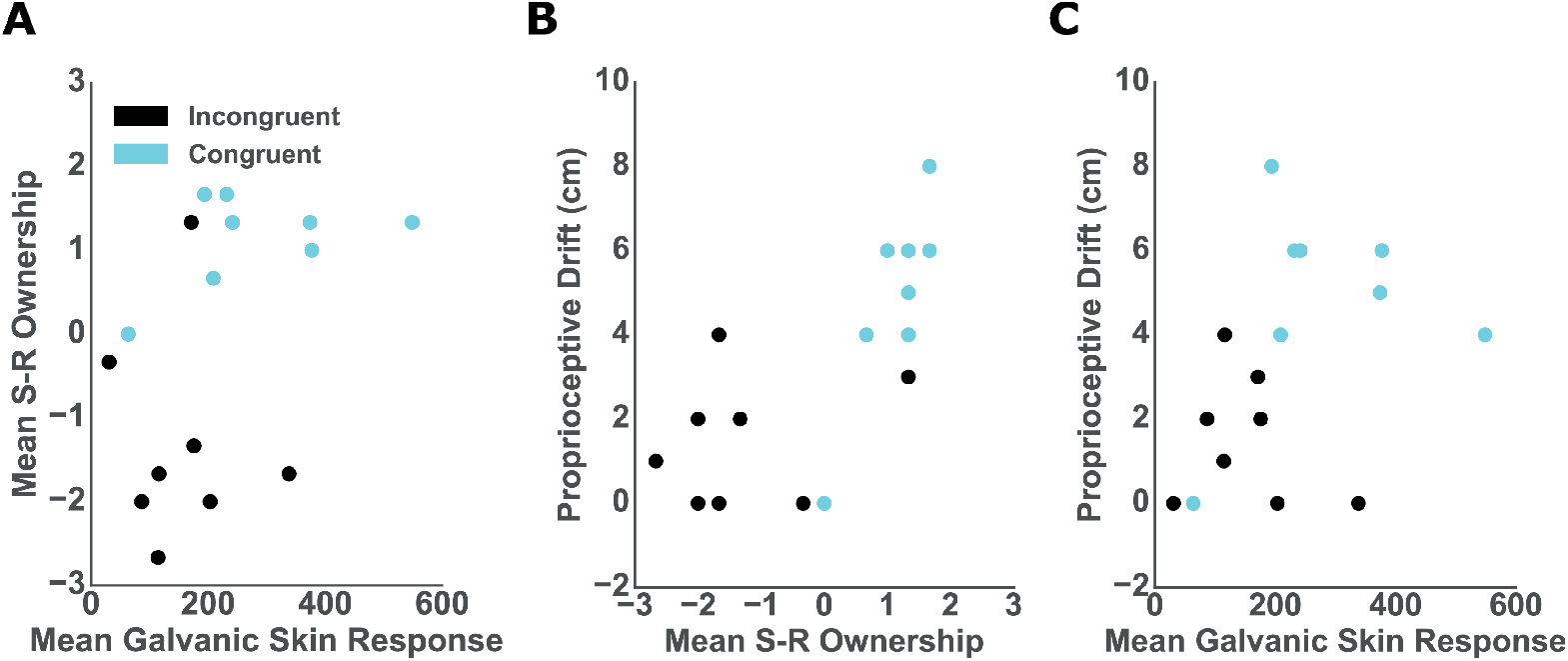
*Correlations* In all graphs dots represent individual participants and colors represent conditions: blue- “C” and black- “I”. (A) Mean GSR 9s post-threatening event and mean self-reported ownership. (B) Mean self-reported ownership and PD score. (C) Mean GSR 9s post-threatening event and and PD score.

### Body Ownership

The analysis revealed that the post-threatening stimulus GSR was significantly higher in “C” (*µ* = 42.54, *sd* = 33.98) than in “I” (*µ* = 29.67, *sd* = 26.82) *p <* 0.001, *d* = 0.42 (Fig 2 F, G). Similarly, we found a difference in the proprioceptive drift between “C” (*µ* = 4.88, *sd* = 2.36) and “I” (*µ* = 1.5, *sd* = 1.51) such that the errors in “C” were significantly higher than in “I” (*t* = 3.4, *p* = 0.004, *d* = 1.7) (Fig 2 H). We further report a statistically significant difference in the self-reported experience of ownership between the two conditions (*t* = 4.97, *p <* 0.001, *d* = 2.5). The ownership ratings in “C” (*µ* = 1.13, *sd* = 0.56) were greater than in “I” (*µ* = *−*1.3, *sd* = 1.25). We found no difference between “C” (*µ* = *−*1.33, *sd* = 1.46) and “I” (*µ* = *−*1.3, *sd* = 1.25) for the three control items (*t* = *−*1.79, *p* = 1.38). We later analyzed questions related to agency. The results showed differences neither for the control questions (*t* = 0.22, *p* = 0.82) between “C” (*µ* = *−*1.67, *sd* = 1.49) and “I” (*µ* = *−*1.83, *sd* = 1.48) nor for the experimental ones, “C” (*µ* = 1.5, *sd* = 1.13) and “I” (*µ* = 1.33, *sd* = 1.48). In both conditions participants experienced high agency during the experiment.

### Correlations

We assessed the relationship between the objective, subjective and behavioral ownership measures and, per each participant in both conditions, we computed: (1) mean GSR from nine seconds post-threat, (2) mean of the three ownership questions; and (3) baseline-subtracted proprioceptive drift. The Spearman rank-order correlation between post-threat GSR and self-reported ownership was close to significance (*r* = 0.47; *p* = 0.06) (Fig 3 A). However, we report high and significant positive correlation between the proprioceptive drift and self-reported ownership (*r* = 0.75; *p <* 0.001) (Fig 3 B) as well as between the post-threat GSR and proprioceptive drift (*r* = 0.52; *p <* 0.03) (Fig 3 C).

## Discussion

In the present study, we asked the question of weather Body Ownership (BO) depends on the consistency of FMs driving goal-oriented action and the proximal signals received from the body (Task-Relevant Proximal Cues, TRPC) as well as distal ones triggered by events outside of the body (Task-Relevant Distal Cues, TRDC). In particular, we investigated the influence of TRDC on performance and BO using an embodied, VR-based, goal-oriented task where action outcomes were signaled by distinct TRDC in the auditory domain. We hypothesized that (in)congruency of TRDC would affect both performance and BO. Our results confirm this hypothesis and demonstrate that both are reduced when the TRDCs are incongruent.

The plasticity of BO relative to the congruency of proximal interoceptive and exteroceptive (both efferent and afferent) signals is well accepted [15, 22, 41–43]. In particular, using the standard method of ownership manipulation (RHI) [5] or VR-based visuomotor (a)synchrony paradigms, where BO is established by either externally-generated visuotactile or self-generated visuomotor synchrony, neurophysiological and behavioral studies have demonstrated that BO results from bottom-up and top-down multisensory matching [15, 42]. Hence, changes in self-specific priors (incorporating a fake body part into self-representation) are driven by statistical correlations when conflicts between visual, tactile, and proprioceptive inputs tend to zero [5]. Current theoretical action–perception framework, grounded in predictions, interprets this underlying mechanism driving BO as minimization of Sensory Prediction Errors (SPE), which during RHI leads to bias in a multisensory generative model [16, 22, 43]. Similar interpretations have been proposed in contexts when the sensory cues are self-generated suggesting that, when acting, BO might result from a computation of an internal Forward Model (FM) [23].

Crucially, however, the discussed paradigms [18–20] do not include manipulation of the environment, which constrains our understanding of BO to self- and externally-generated proximal cues (TRPCs) exclusively. To study whether the plasticity of BO depends on the consistencies of internal FMs and therefore not only TRPC but also TRDC following volitional goal-oriented behavior, we used a paradigm which required the participants to perform actions that trigger TRDC and manipulated their congruency. We predicted that BO scores might be lower in the condition where the cues do not match sensory predictions. Results from all the ownership measures (Fig 2 Lower panel: body Ownership), including GSR, PD and the questionnaire support that TRDC which pertain to the task and violate predictions about the action outcomes compromise BO. Specifically, we found that the BO scores were significantly higher in the congruent compared to the incongruent condition in all analyses. Subsequent correlations between the proposed measures (Fig 3) further confirmed the consistency of the obtained results within three dimensions of ownership quantification including physiological response, behavioral proprioceptive recalibration, and a conscious report [40].

Similarly to the asynchronous stroking condition in the standard RHI paradigm [5] or visuomotor mismatch [19], here we interpret the obtained low-ownership outcome in the incongruent condition (Fig 2 Upper panel: body Ownership) as a consequence of high sensory SPE of FMs or the efference copy [16, 28]. In our case, however, the sensory conflicts are driven by a discrepancy between the predicted and actual distal auditory signals which do not pertain to the body. In particular, the manipulation of TRDC might have reflected on the errors of the FMs which drive both performance and BO. This could suggest TRDC might influence TRPC establishing a feedback loop such that any (in)congruent relationship between them which pertain to the goal will affect BO and even define the boundaries of the embodied self. To the best of our knowledge, this result proposes for the first time that BO might be driven by bottom-up integration and top-down prediction of not only proximal sensory cues deriving from the body but also distal ones occurring outside of the body. This would support results from other studies showing that BO is coupled to the motor systems [37, 44] and that it might depend on FMs. Consistent with other studies [20], the differences in the perceived BO were not influenced by a lack of agency (Fig 2 J). In particular, the participants experienced control over the virtual hand in both groups, probably due to the congruent mapping of the proximal cues. The visual feedback of the movement of the arm always followed the desired trajectory, which is one of the crucial questions addressed in the standard self-reported agency assessment [40].

Crucially, at the current stage, two questions still remain open. First, how can the integration of TRDC and TRPC occur in the service of BO? For example, a recent Hierarchical Sensory Predictive Control (HSPC) theory proposes a cascade of purely sensory predictions which mirror the causal sequence of the perceptual events preceding an event [35]. This control architecture acquires internal models of the environment through a hierarchy of sensory predictions from visual (distal, TRDC) to proprioceptive and vestibular modalities (proximal, TRPC). If BO and motor control share the same FMs which comprise both TRDC and TRPC during a goal-oriented task, BO might be realized through a similar cascade of sensory predictions. Second, if BO depends on the consistency of internal models, and therefore on the accuracy of sensory predictions, could task-irrelevant signals contribute to BO? Changing the rules of the environment and investigating BO and performance when action-independent (task-irrelevant) sensory predictions are violated would shed light on the nature of sensory signals relevant for the processing of self and their underlying mechanisms (ie. generative and forward models) [16, 22, 45].

What is the role of TRDC in goal-oriented behavior? Our results demonstrate that performance, measured as overall scores (Fig 2 A) and directional errors (Fig 2 B, C), was significantly hampered in the incongruent compared to the congruent condition. Importantly, these results did not depend on a difference in RTs (Fig 2 E) suggesting no influence of possible attentional biases (ie. distractions) in either of the groups. On the one hand, this outcome might be interpreted from a computational motor control perspective. The reported differences in performance between the conditions could have been influenced by the discrepancies between the efference copies of distal events and the actual action outcomes. Indeed, results from motor control studies support the notion that learning (progressive reduction of error) depends on both proximal and distal sensory prediction errors that allow for adjustments and anticipation of possible perturbations deriving from the body and environment [30, 35, 46–49]. As a result, inputs from all the sensory modalities are transformed into error signals while an action is being executed, updating the forward model and, consequently, future behavior [29, 31, 50, 51]. In our experiment, the directionality of the error indicated by the spatial distribution of the sound (left or right) as well as its temporal characteristics could both be interpreted as error signals which supervise corrective commands. Thus TRDC in the incongruent condition might have influenced performance, which, in turn, affected body ownership. For example, clinical studies provide evidence that patients suffering from hemiparesis, whose motor function is reduced due to stroke, progressively stop using the paretic limb: the so-called learned non-use phenomenon [52]. In this and other neurological cases, permanent lack of use (low performance) often causes disturbances in the sense of ownership and agency [53] supporting a hypothesis that there might be a causal effect between performance and body ownership. The present design, however, which includes three types of sensory manipulations pseudorandomly distributed within each block, does not allow us to disambiguate between the specific contribution of each of TRDC. A systematic study on the influence of individual sensory signals, including the three manipulations, would help to better understand the mechanisms accounting for low-performance scores in the incongruent condition.

An alternative interpretation comes from the body ownership literature. Several studies propose that BO is coupled to the motor system such that it updates the sensory representation of the body and provides inputs to the Forward Model (FM). The FM, in turn, generates and updates predictions relative to both the body and the environment during voluntary actions [44], reinforcing the history of sensorimotor contingencies. In particular, we find evidence that BO is involved in generating body-specific predictions about the sensory consequences of voluntary actions thus determining somatosensory attenuation [44]. This outcome is consistent with another study using standard RHI in VR [37], which shows that the degree of ownership correlates with motor performance in a decision-making task. Contrary to the previous discussion, in this case, ownership would have a modulatory effect on performance.

At the current stage, we cannot disambiguate between the two alternative hypotheses and determine whether TRDC influence BO and performance in parallel or sequentially and what is the directionality. We demonstrate, however, that both depend on TRDC, which supports that both depend on the consistency of FMs driving goal-oriented action and both proximal and distal cues which this action generates pertaining to both the body and the environment [16, 22]. We expect that this outcome will allow for the advancement of our understanding of the mechanisms underlying body ownership.

Finally, the reported finding might find applications in fields such as motor training simulators and rehabilitation. For instance, virtual reality-based treatments of post-stroke motor disorders [54–57] might benefit from a design of reliable and spatiotemporally congruent environments which increase body ownership and self-representation possibly impacting recovery. Further clinical studies should evaluate the same principle in rehabilitation protocols for ownership disturbances following acquired brain lesions including neglect [58], anosognosia for hemiplegia [59] or somatoparaphrenia [60].

## Author Contributions

K.G., L.U. and B.R.B. designed the protocol, K.G and L.U. conceived the experiment and L.U. conducted the experiments, K.G. and L.U. analyzed the results, K.G., L.U., B.R.B. and P.F.M.J.V. wrote the manuscript. P.F.M.J.V. initiated and supervised the research. All authors reviewed and approved the manuscript.

